# Musical expertise shapes visual-melodic memory integration

**DOI:** 10.1101/2022.02.03.478977

**Authors:** Martina Hoffmann, Alexander Schmidt, Christoph J. Ploner

## Abstract

Music can act as a powerful mnemonic device that can elicit vivid episodic memories. However, how musical information is integrated with non-musical information is largely unknown. Here, we investigated whether and how musical expertise modulates binding of melodies and visual information into integrated memory representations. We reasoned that the significant mnemonic demands of musicianship might alter the underlying integration process and reveal mechanisms by which music promotes retrieval of non-musical memories. Professional musicians and musical laypersons learned overlapping pairs of everyday objects and melodies (AB- and BC-pairs, object-melody and melody-object pairs). Participants were then tested for memory of studied pairs (direct trials) and for inferential AC-decisions (indirect trials). Although musicians showed a higher overall performance than non-musicians, both groups performed well above chance level in both trial types. Non-musicians reacted faster in indirect compared to direct trials, whereas the reverse pattern was found in musicians. Differential correlations of trial type performance between groups further suggested that non-musicians efficiently formed integrated ABC-triplets already during the encoding phase of the task, while musicians separately memorized AB- and BC-pairs and recombined them at retrieval for AC-decisions. Our results suggest that integrative encoding is a default mechanism for integration of musical and non-musical stimuli that works with great efficacy even in musically untrained subjects and may contribute to the everyday experience of music-evoked episodic memories. By contrast, recombination at retrieval seems to be an advanced strategy for memory integration that critically depends on an expert ability to maintain and discriminate musical stimuli across extended memory delays.

## Introduction

Music and memory are intimately related. Listening to specific songs or melodies can evoke a multitude of vivid episodic and autobiographical memories. Previous research identified the association of music with memories as a main motivation for music listening^1^. Recent empirical studies confirmed that people across all age groups use music to reflect on the past and deliberately evoke personal memories^2,3^. Moreover, connection with autobiographic memory episodes is a major mechanism for the experience of emotions when listening to music^4–6^ and might partially explain hippocampal involvement in music-evoked emotions^7^. Thus, musical information can be associated with non-musical information, such as semantic and episodic memories, emotions and motor programs^6,8,9^.

In addition to music listening, active music making also critically depends on learning and memory^10–12^. In classical music, for example, it is common practice for musicians to memorize entire concertos or other musical pieces and perform them without sheet music. In jazz, musicians usually learn a large repertoire of tunes and know their musical structure, melodies and harmonic progressions by heart. Moreover, musical training has been shown to foster active and controlled learning strategies^9,13^. Music making, in particular at a professional level, therefore is accompanied by changes in brain areas that are involved in learning and memory. For instance, gray-matter volumes in the hippocampus differ between musicians and non-musicians and increase with the amount of musical expertise^14^. In addition, stronger hippocampal activation was found in musicians compared to non-musicians in a musical familiarity task which might indicate that musicians have specific memory abilities^15^.

So far, it has not been investigated whether the abovementioned changes transfer on musicians’ abilities to link musical information with non-musical memories. It is currently still a matter of debate whether and how musical expertise modulates memory performance and whether functional and structural brain changes in musicians reflect qualitative differences to non-musicians. In a recent meta-analysis of the effects of musical training on non-musical cognitive and academic skills in children and adolescents, there was no effect of musical training on any outcome, including memory, after controlling for quality of studies^16^. Another meta-analysis on young adults only included studies with memory tasks and found that musicians performed better than non-musicians in short-term memory, working memory and, to a lesser extent, long-term memory tasks^13^. However, the memory advantage of musicians was at least partially domain-specific, with large effect sizes for tonal stimuli, moderate for verbal stimuli and small or null for visuospatial information.

Here, we investigated how professional musicians and non-musicians use music to bind distinct non-musical information into overlapping visual-melodic representations. The process of integrating memories that are related but were not experienced together is called memory integration^17,18^. This cognitive faculty is a major prerequisite for building networks of interrelated memory items and for organizing knowledge^19,20^. Memory integration moreover plays an important role in various non-mnemonic cognitive processes, such as decision making, spatial navigation, acquisition of memory schema, creativity, imagination and concept learning^17–20^. We compared professional musicians and non-musicians by using a visual-melodic variant of an associative inference task^21,22^, a paradigm that is frequently used to study memory integration. Participants were required to memorize overlapping melody-object pairs and to form an integrated memory of these associations. We thus aimed to study the basic mechanisms that underlie the connection of non-musical memories with musical information. We further aimed at revealing possible differences in mnemonic strategies between musicians and non-musicians that might relate to the abovementioned structural and functional brain changes in musicians.

## Methods

### Participants

A total of 60 participants was recruited for the study, 30 professional musicians and 30 non-musicians (Table 1). The professional musicians either studied at a music university or music school or had completed their studies and worked as instrumental teachers, freelance and orchestra musicians. All musicians were instrumental musicians (string instruments n = 12; keyboard instruments n = 6; woodwind instruments n = 5; brass instruments n = 3; plucking instruments n = 4). The non-musician group consisted of 30 participants without or with minimal extracurricular musical activity. Six of these participants reported that they had played or tried a musical instrument or had sung in a school choir for 6 months to 2.5 years. However, musical activity was abandoned at least ten years prior to study participation. Non-musicians were recruited from staff and students of the Charité – Universitätsmedizin Berlin and from other Berlin universities.

**Table 1.** Demographics and musical activity of the musician and non-musician groups. Values are given as means with standard deviations or as frequencies. χ^2^ = chi-square test; y = years; h = hours; W = Wilcoxon rank-sum test; LPS = Leistungsprüfsystem; MBEA = Montreal Battery of Evaluation of Amusia; ^a^ Average hours of daily music listening during the last 12 months; ^b^ Average number of attended concerts or music events during the last 12 months; ^c^ Average practice time per week across all decades of musical activity; ^d^ Average practice time per week during the last 12 months

|  | Musicians<br>(n = 30) | Non-musicians<br>(n = 30) | Test statistic and p<br>value |
| --- | --- | --- | --- |
| Sex (female / male) | 14/16 | 14/16 | $\chi^2(1) = 0$, $p = 1.00$ |
| Age (y) | 25.40 ± 5.87 | 26.37 ± 4.78 | W = 377.5,<br>$p = 0.286$ |
| Years of education (y) | 15.37 ± 2.34 | 15.82 ± 1.66 | W = 404, $p = 0.498$ |
| Reasoning - LPS Subtest<br>No. 3 (T Score) | 61.83 ± 5.94 | 59.67 ± 6.81 | W = 537.5,<br>$p = 0.185$ |
| MBEA Scale subtest (%) | 94.89 ± 5.98 | 87.67 ± 7.12 | W = 727.5,<br>$p < 0.001$ |
| Daily music listening<sup>a</sup> (h) | 1.53 ± 1.17 | 1.89 ± 1.64 | W = 390.5,<br>$p = 0.501$ |
| Concert visits<sup>b</sup> | 27.83 ± 28.61 | 6.80 ± 16.25 | W = 785.5,<br>$p < 0.001$ |
| Age of first instrumental<br>practice (y) | 5.67 ± 1.99 | - | |
| Total years of<br>instrumental practice (y) | 19.05 ± 5.54 | 0.22 ± 0.59 | |
| Accumulated instrumental<br>practice time (h) | 17448 ± 9005 | 14 ± 44 | |
| General average weekly<br>practice time<sup>c</sup> (h) | 16.02 ± 7.62 | 0.22 ± 0.44 | |
| Current average weekly<br>practice time<sup>d</sup> (h) | 24.05 ± 12.11 | - | - |
| Absolute pitch (yes/no) | 9/21 | - | - |

No participant reported a history of neurological or psychiatric diseases, hearing deficits or significant visual impairments. The musician and non-musician groups were matched for sex, age and educational level (Table 1). Both groups were comparable in terms of non-verbal intelligence as measured with a logical reasoning task (Subtest 3 of the test battery Leistungsprüfsystem LPS)^23^. The Scale Subtest of the Montreal Battery of Evaluation of Amusia (MBEA)^24^ was used to screen for amusia and assess basic music perceptual abilities. Although musicians outperformed non-musicians in this test, all non-musician participants scored within the normal range. All participants gave written informed consent before participation in the study. The study was approved by the local Ethics Committee of the Charité – Universitätsmedizin Berlin and was conducted in conformity with the Declaration of Helsinki.

### Musical associative inference task

We used a musical adaption of a visual associative inference task that has previously been used to study memory integration in behavioral and fMRI studies^21,22,25–27^ (Figure1). In the musical variant, participants learned overlapping pairs of objects and melodies (object-melody and melody-object pairs; i.e. AB- and BC-pairs) and non-overlapping object-melody associations (DE-pairs). After a memory delay, participants were tested both on studied AB-, BC- and DE-pairs (‘direct trials’) and on inferential AC-associations (‘indirect trials’).

**Figure 1.**
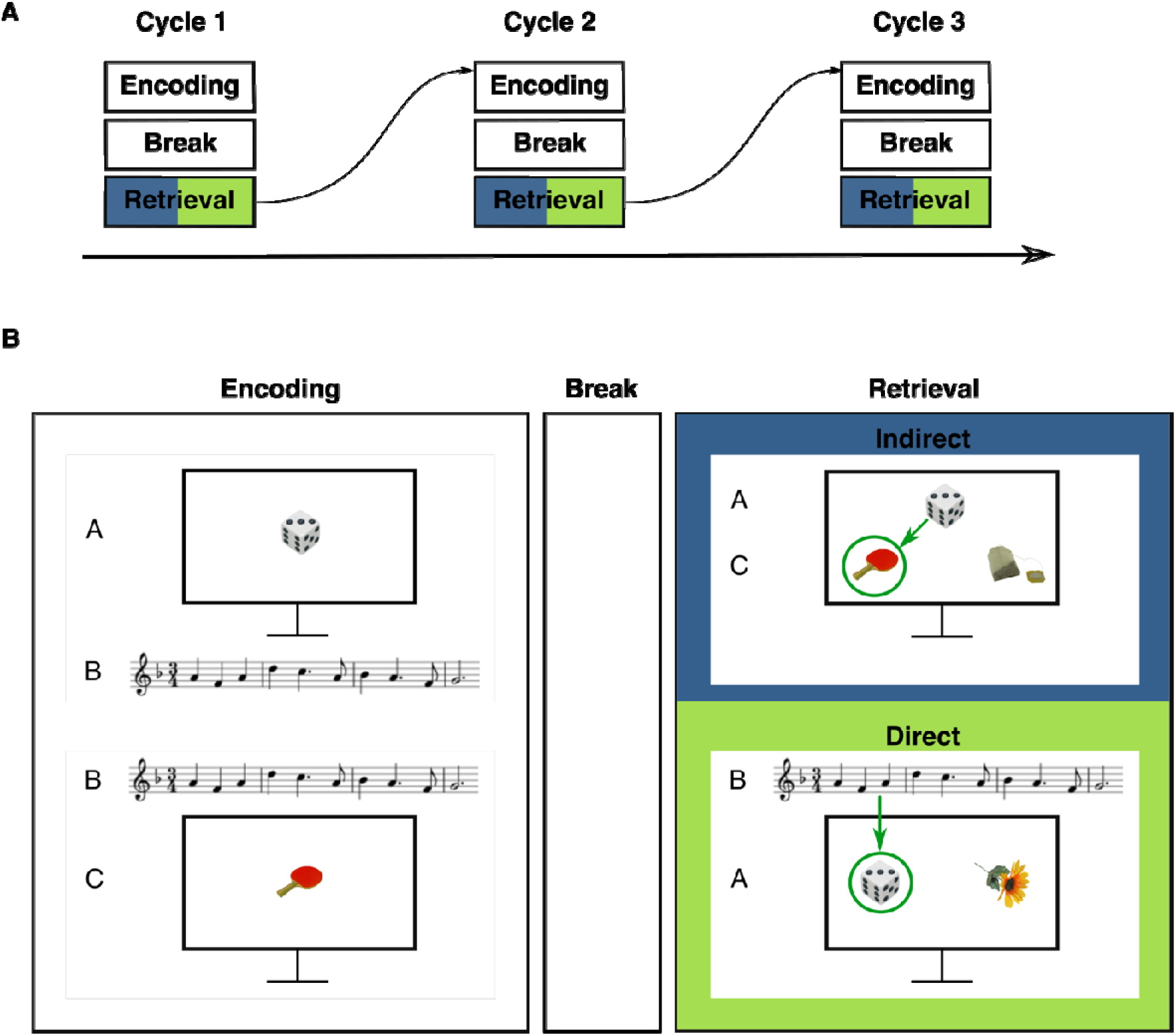
Procedure and example stimuli of the musical associative inference task. (A) Task structure: The experiment consisted of three alternating encoding and retrieval blocks that were separated by a break. (B) Example stimuli of encoding and retrieval blocks. During the encoding block, participants studied overlapping pairs of objects and melodies (AB-/BC-pairs) and non-overlapping DE-pairs (not shown). Note the overlapping melody of AB-/BC-trials. At retrieval, participants were tested on studied direct trials (AB-, BC-, DE-pairs) and on indirect trials (inferential AC-pairs). Green arrows indicate the correct choice.

#### Stimuli

Visual stimuli were taken from the Bank of Standardized Stimuli (BOSS Phase II)^28^ and consisted of 331 colored images of everyday objects (e.g. tools, food, clothes, toys etc.). Musical stimuli involved 43 lesser known melodies played in a piano voice, without orchestration and lyrics, even if the original piece included lyrics. Melodies were taken from various genres, such as classical music, jazz, folk songs (from non-German speaking countries) or themes from older TV series or movies (see Supplementary Table 1). The melodies had a mean duration of 7 seconds (Range: 5-10 sec).

#### Procedure

The experiment consisted of three cycles with one encoding and retrieval block in each cycle. Encoding and retrieval blocks were separated by a delay of five minutes (Figure 1). Each encoding block consisted of 18 trials with melody-object pairs (six AB-, six BC- and six DE-trials). Participants were presented an object (A) on the computer screen and a melody (B) at the same time. Two to four trials later, the same melody (B) was played again, but was now paired with another object (C). A- and C-stimuli were always objects and the overlapping B-stimulus always a melody. In addition, non-overlapping DE-trials were presented, consisting of one object (D-stimulus) and one melody (E-stimulus). These stimuli did not overlap with other trials of the encoding blocks. DE-trials were included in the design to establish a minimum distance between AB- and BC-trials and to increase uncertainty about occurrence and timing of BC-trials. The combination of objects and melodies was trial-unique and pseudo-randomized for each participant. Within each encoding block, the order of the trials was pseudo-randomized using the program Mix^29^. AB-pairs were always presented before their corresponding BC-pairs. DE-trials were intermixed with AB- and BC-trials. Presentation time of each pair was determined by the length of the respective melody. In order to ensure that participants focused on the presented stimuli, participants were asked how they liked the melodies after each trial. Responses were given on a five-point Likert scale (1 = not at all, 5 = very much). Trials were terminated after a response was given. The inter-trial interval was 5 seconds.

After each encoding block, a break of five minutes was inserted. Then, the corresponding retrieval block was started. During retrieval, testing of AC-trials always preceded the retrieval of direct associations. Thus, relearning of AB- and BC-trials before testing of AC-pairs was prevented. In each cycle, participants were tested on six AC-trials. At the top of the screen, one A-stimulus (i.e. an object) was presented (Figure 1). Two C-stimuli were shown below, one representing the target object and one a foil object. Participants had to decide which of the two C-stimuli at the bottom had previously been presented with the same melody as the A-stimulus. Participants indicated their choice via button press. For retrieval of direct associations, all 18 AB-, BC- and DE-trials of the respective cycle were tested. In the middle of the computer screen, two objects (i.e. either two A-, C- or D-stimuli) were shown. At the same time, a melody was played (either a B- or E-stimulus). Participants had to indicate by button press which of the two objects had previously been paired with the melody (Figure 1). The order both of AC- and direct trials was randomized. The presentation of the stimuli was terminated by key press of the participants. To avoid differences in familiarity of target and foil stimuli, all foils were taken from different pairs of the same cycle. In each retrieval block, tested pairs were always from the encoding block of the same cycle. No objects or melodies from other than the respective cycle were presented.

The experiment was performed using Presentation◻ software (Version 18.1, Neurobehavioral Systems, Inc. Berkeley, CA) and took place in a quiet room. Musical stimuli were presented via external speakers and participants could adapt the volume to their needs. Prior to the experiment, participants were instructed about the task with example stimuli and received a short training with a small number of trials. Melodies and objects of the training session were not included in the proper experiment. The experimenter ensured full comprehension of instructions before the experiment was started.

### Assessment of musical activity

Indices of musical activity were assessed using a short questionnaire (MusA)^30^. This questionnaire covers both music reception (i.e. music listening, concert attendance) and active musical practice (i.e. instrument group, years of musical activity, weekly practice time). The weekly average time of musical practice was assessed for each age decade (i.e. 0-10 years, 11-20 years, 21-30 years etc.). Additionally, weekly average time of music making during the last 12 months and total years of instrumental practice were measured. The variables assessed via the MusA were used to calculate further indices of musical activity. The general average practice time across all decades was determined by calculating the mean of the weekly average time of playing music for each age decade. The cumulative practice time on the instrument was calculated by combining total years of instrument playing with weekly practice times. In addition, we assessed musical activity variables that were not covered by the questionnaire by using a short personal interview (age of first instrumental practice, played instruments, absolute pitch).

### Data analysis

For the musical associative inference task, we analyzed accuracy, i.e. the percentage of correct responses for each trial type (AC-trials and direct trials, i.e. AB-, BC- and DE-trials) in each participant. We further analyzed reaction times (RTs) of correctly answered trials for each trial type. Medians were used to describe individual average RTs for each trial type. Due to the limited number of trials per cycle and trial type, data were averaged across cycles.

Since most of the variables of interest were not normally distributed, a non-parametrical statistical approach was used throughout. Analyses were performed using R Studio^31^ (version 3.6.3). Accuracy and RTs were analyzed with a repeated measures design for non-normal data using the package MANOVA.RM^32,33^. With this package, robust test statistics can be calculated if the assumptions of parametric approaches (i.e. normal distribution, equal covariances) are violated. We calculated Wald-type statistics (WTS) with permuted p-values to account for non-normal data distribution. Significant interactions were followed by pairwise comparisons. For post-hoc comparison of within factors (i.e. trial type, response pattern), one-way repeated measure ANOVAs were performed with the RM function of the MANOVA.RM package. For post-hoc analysis of group differences, we used the package GFD^34^ to calculate WTS combined with a permutation procedure for p-values. The Bonferroni-Holm correction^35^ was used to adjust for multiple comparisons in the post-hoc analysis. For comparison of demographic data and musical activity variables across groups, Wilcoxon rank-sum tests were calculated. Kendall’s τ was calculated for determining correlations (e.g. between accuracy of different trial types and between accuracy and musical activity). The significance level was set at p < .05.

## Results

### Accuracy

We first analyzed differences in accuracy between direct trials with an overlapping melody (i.e. AB- and BC-trials) and direct trials without melody repetition (i.e. DE-trials) and conducted a repeated measures ANOVA for non-normal data with group (musicians, non-musicians) as between-factor and direct trial type (AB-/BC-trials, DE-trials). Since the main effect of direct trial types (WTS(1) < 1, p = 0.393) or the interaction between group and direct trial type (WTS(1) = 3.23, p = 0.076) was not significant, we pooled all direct trials (i.e. AB-, BC- and DE-trials) for further analysis, similar to previous studies^21,26^.

Data of the two groups and trial types are plotted in Figure 2. In a first step, we checked whether both groups performed above chance level (i.e. 50% correct answers) in indirect and direct trials using a Wilcoxon signed rank test. In both trial types, performance was significantly above chance level in musicians (indirect trials: M = 79.44%, SD = 17.08%, W = 457.5, p < 0.001; direct trials: M = 84.44%, SD = 9.79%, W = 465, p < 0.001) and non-musicians (indirect trials: M = 71.29%, SD = 16.38%, W = 367, p < 0.001; direct trials: M = 73.83%, SD = 9.61%, W = 465, p < 0.001).

**Figure 2.**
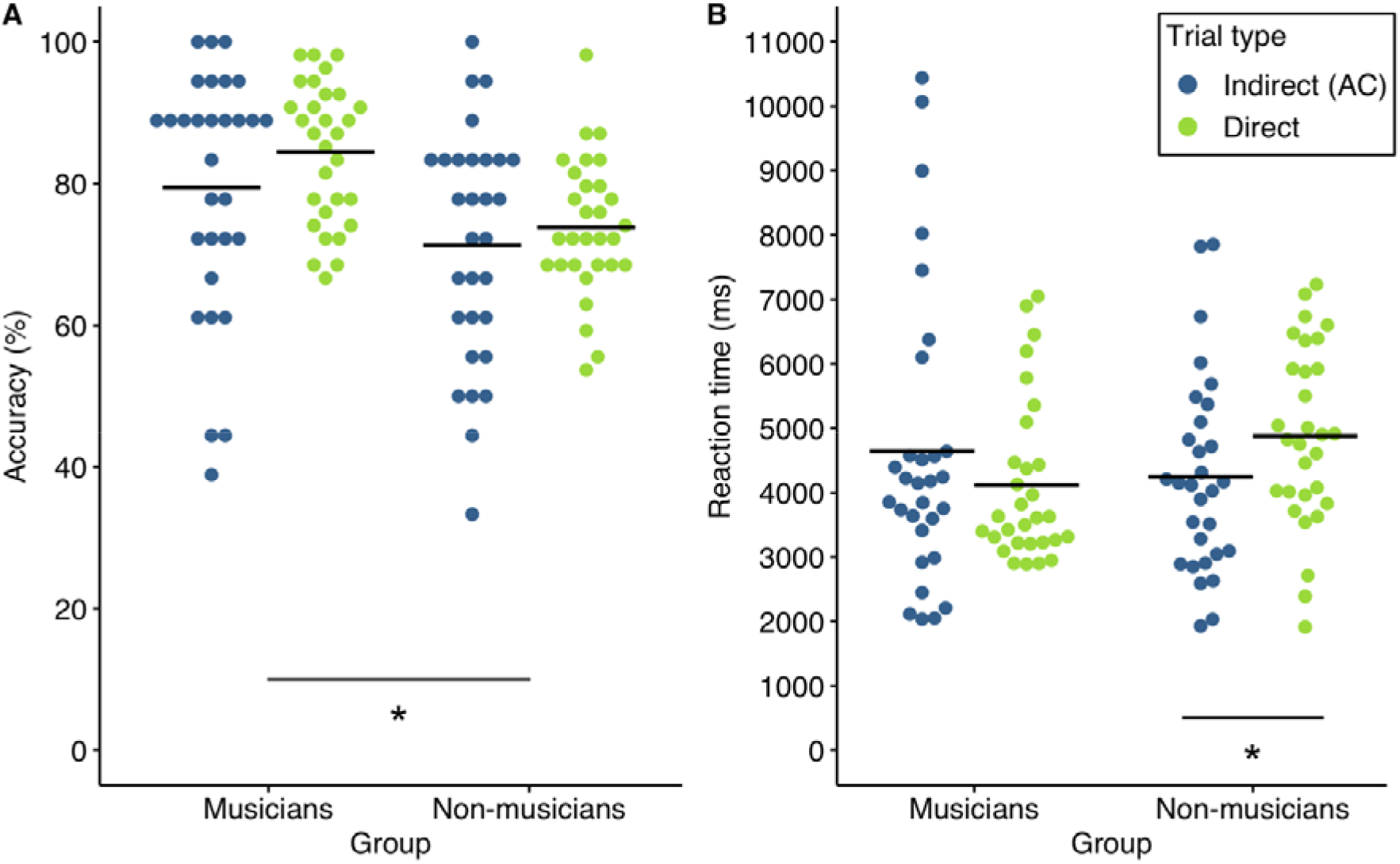
Accuracy and reaction times of both groups. (A) Accuracy in indirect (blue) and direct (green) trial types in musicians and non-musicians. There was a significant group effect (* p < 0.05) (B) Reaction times in indirect (blue) and direct (green) trial types in musicians and non-musicians. There was a significant interaction of group x trial type. The asterisk denotes the significant pairwise comparisons (*p < 0.05 after Bonferroni-Holm correction). Solid lines represent the respective mean.

Accuracy was then analyzed using a repeated measures ANOVA for non-normal data with group (musicians, non-musicians) as between-factor and trial type (indirect, direct) as within-factor. There was a significant group difference (WTS(1) = 10.49, p = 0.002). Averaged across trial types, musicians (M = 81.94%, SD = 12.12%) performed superior to non-musicians (M = 72.56%, SD = 10.25%). There was no significant effect of trial type (WTS(1) = 3.47, p = 0.072) or interaction of trial type and group (WTS(1) <1, p = 0.545). Although musicians outperformed non-musicians in both trial types, non-musicians were also able to efficiently used melodies (i.e. B-stimuli) to associate non-musical information.

### Reaction Times

Data of the two groups and trial types are plotted in Figure 2. For analysis of RTs of correctly answered trials, a repeated measures ANOVA with group (musicians, non-musicians) as between factor and trial type (indirect, direct) as within factor was calculated. There was no significant main effect of group (WTS(1) < 1, p = 0.623) or trial type (WTS(1) < 1, p = 0.828), indicating that non-musicians were generally as fast as musicians in retrieving associations between musical and non-musical information and that RTs were not generally shorter in one of the trial types. The interaction effect of group and trial type however was significant (WTS(1) = 7.1, p = 0.009). We thus compared the respective levels of the factors trial type and group (corrected for four pairwise comparisons). Post-hoc tests showed trial type differences for non-musicians (WTS(1) = 9.34, p = 0.016). RTs were significantly shorter in indirect trials (M = 4249ms, SD = 1526ms) compared to direct trials (M = 4882ms, SD = 1392ms). The musician group showed the opposite pattern with longer RTs in indirect trials (M = 4652ms, SD = 2276ms) compared to direct trials (M = 4118ms, SD = 1249ms). However, this difference did not achieve significance in the post-hoc test (WTS(1) = 1.917, p = 0.362). Post-hoc tests between the two groups did not show significant differences for indirect (WTS(1) < 1, p = 0.427) or direct trials (WTS(1) = 5.02, p = 0.081) after correction for multiple comparisons. We reasoned that the different RT patterns might reflect different strategies for memory integration in the two groups. Shorter RTs in indirect trials compared to direct trials in the non-musician group might suggest that non-musicians build ABC-associations already during encoding of BC-stimuli with the preceding AB-stimuli and thus react faster in later indirect AC-decisions. By contrast, the RT pattern in the musician group might suggest a more retrieval-based strategy with musicians separately memorizing AB- and BC-pairs and then combining them at retrieval for AC-decisions.

### Accuracy of corresponding direct and indirect trials

To test our hypothesis of different memory integration strategies underlying the distinct RT patterns between groups, we further investigated how musicians and non-musicians used direct AB- and BC-trials for indirect AC-decisions. We reasoned that a mainly retrieval-based strategy for associative inference necessitates correct memory of AB- and BC-pairs for correct AC-decisions at retrieval.

Conversely, if AC-pairs are already formed at encoding (i.e. as soon as a BC-pair is presented) and are remembered until retrieval, AC-decisions may be less dependent on correct memory of AB- and BC-pairs. We therefore analyzed the performance of the corresponding trials of one triplet (AC-, AB-, BC-pairs). In a first step, we took AC-trials that were correctly answered and checked whether the corresponding AB- and BC-trials were correct or incorrect, resulting in four different response patterns. Relative percentages for each participant were then calculated by dividing the number of each response pattern by the respective number of correctly answered AC-trials. Percentages of the trials in which AB or BC or both were incorrectly answered were summed up, resulting in two response patterns: (1) AB- and BC-trials were correct, indicating the percentage of all correct AC-trials in which the corresponding AB- and BC-trials were also correctly answered (i.e. AB-, BC- and AC-trials of one overlapping ABC-triplet were correctly answered). (2) AB- and/or BC-trials were incorrect, indicating the percentage of all correct AC-trials in which the corresponding AB or BC or both were incorrectly answered (i.e. AC-trials were correct, but either AB- or BC- or both trials of the same overlapping ABC-triplet were incorrectly answered). Figure 3 displays data of the two groups and response patterns.

**Figure 3.**
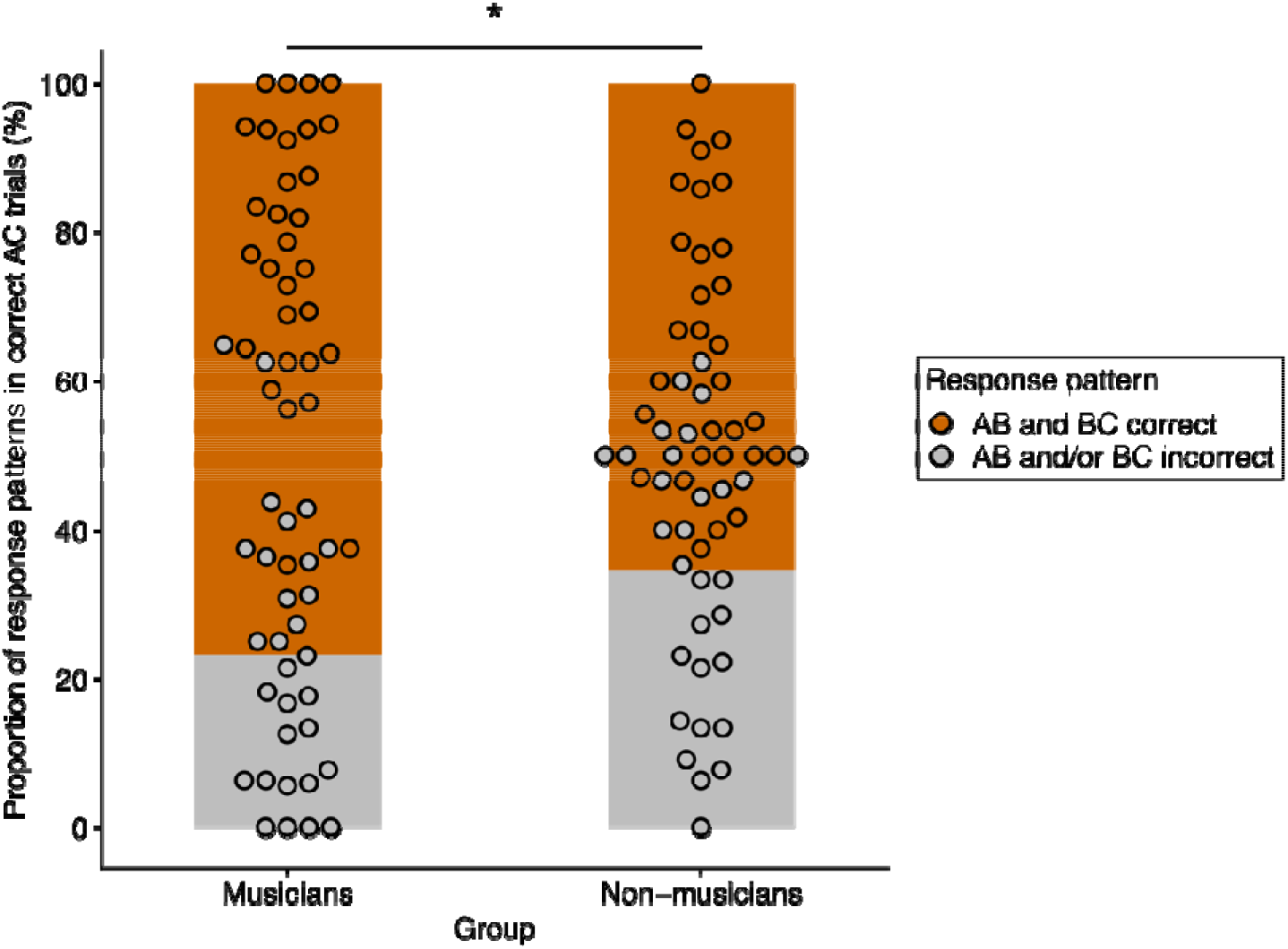
Relative frequencies of the two response patterns in musicians and non-musicians. AB and BC correct (orange) refers to the relative percentage of correctly answered AC-trials in which the corresponding AB- and BC-trials of the same overlapping ABC-triplet were also correctly answered. AB and/or BC incorrect (gray) refers to the relative percentage of correctly answered AC-trials in which the corresponding AB- or BC- or both trials of the same overlapping ABC-triplet were incorrectly answered. There was a significant effect of response patterns and a significant interaction of response pattern and group. The asterisk denotes the significant pairwise comparison of response patterns between groups (*p < 0.05 after Bonferroni-Holm correction).

We then calculated a repeated measures ANOVA for non-normal data with group (musicians, non-musicians) as between-factor and response pattern (AB and BC correct, AB and/or BC incorrect) as within-factor. There was a significant main effect of response pattern (WTS(1) = 83.15, p < 0.001) and a significant interaction effect of group and response pattern (WTS(1) = 6.11, p = 0.016). The main effect of group was not significant (WTS(1) = 1.65, p = 0.21). Post-hoc analysis (corrected for four pairwise comparisons) revealed response pattern differences for both musicians (WTS(1) = 68.12, p = 0.004; AB and BC correct: M = 76.81%, SD = 17.79%; AB and/or BC incorrect: M = 23.19%, SD = 17.79%) and non-musicians (WTS(1) = 21.79, p = 0.004; AB and BC correct: M = 65.37%, SD = 18.04%; AB and/or BC incorrect: M = 34.63%, SD = 18.04%). Not surprisingly, the underlying AB- and BC-trials were correct in the majority of correctly answered AC-trials in both musicians and non-musicians. However, when we compared response patterns between groups, we found significant differences for both response patterns (AB and BC correct: WTS(1) = 6.11, p = 0.035; musicians: M = 76.81%, SD = 17.79%; non-musicians: M = 65.37%, SD = 18.04%; AB and/or BC incorrect: WTS(1) = 6.11, p = 0.035; musicians: M = 23.19%, SD = 17.79%; non-musicians: M = 34.63%, SD = 18.04%). If AC-trials were correct, musicians had a higher percentage of trials in which both the corresponding AB- and BC-pairs were also correct than non-musicians. Non-musicians had a higher percentage of AC-trials in which the corresponding AB- or BC-trials or both were incorrect, suggesting that they could still make correct AC-decisions, even if they did not correctly remember the underlying direct pairs. This analysis suggests that the two groups differed in how they used direct AB- and BC-trials for indirect AC-decisions: musicians mainly based their AC-decisions on knowledge of direct pairs for AC-decisions, whereas non-musicians seemed to use a more encoding-based strategy in which knowledge of direct AB- and BC-trials was less necessary for correct AC-decisions.

### Correlation of accuracy between indirect and direct AB-/BC-trials

We further analyzed the correlational pattern between accuracy in indirect trials (AC) and the overlapping direct trial types (AB, BC) in both groups to better understand the behavioral strategies in the two groups. If musicians indeed based their AC-decisions on knowledge of the underlying direct pairs, a correlation between AC-trials with AB- and BC-trials can be expected. In non-musicians, however, no correlations should be expected if AC-pairs are already formed during encoding and AC-decisions are less dependent on explicit knowledge of AB- and BC-pairs.

In the musician group, correlation analyses revealed significant correlations between AC accuracy and performance in the underlying direct trial types (AC-AB: τ = 0.42, p = 0.0033; AC-BC: τ = 0.3, p = 0.033). No correlation between AC performance and accuracy in AB- or BC-trials was observed in non-musicians (AC-AB: τ = 0.044, p = 0.76; AC-BC: τ = 0.051, p = 0.71). Figure 4 displays the correlation plots for both groups and the respective bivariate correlations. In line with the RT patterns and the analysis of the corresponding direct and indirect trials, the results of the correlation analysis support the hypothesis of different behavioral strategies for memory integration in musicians and non-musicians. Musicians relied on memory of underlying AB- and BC-pairs when deciding on AC-trials, while non-musicians tended to form ABC-representations during encoding for later AC-decisions. AC-decisions in this group were thus less dependent on correct memory of direct trials.

**Figure 4.**
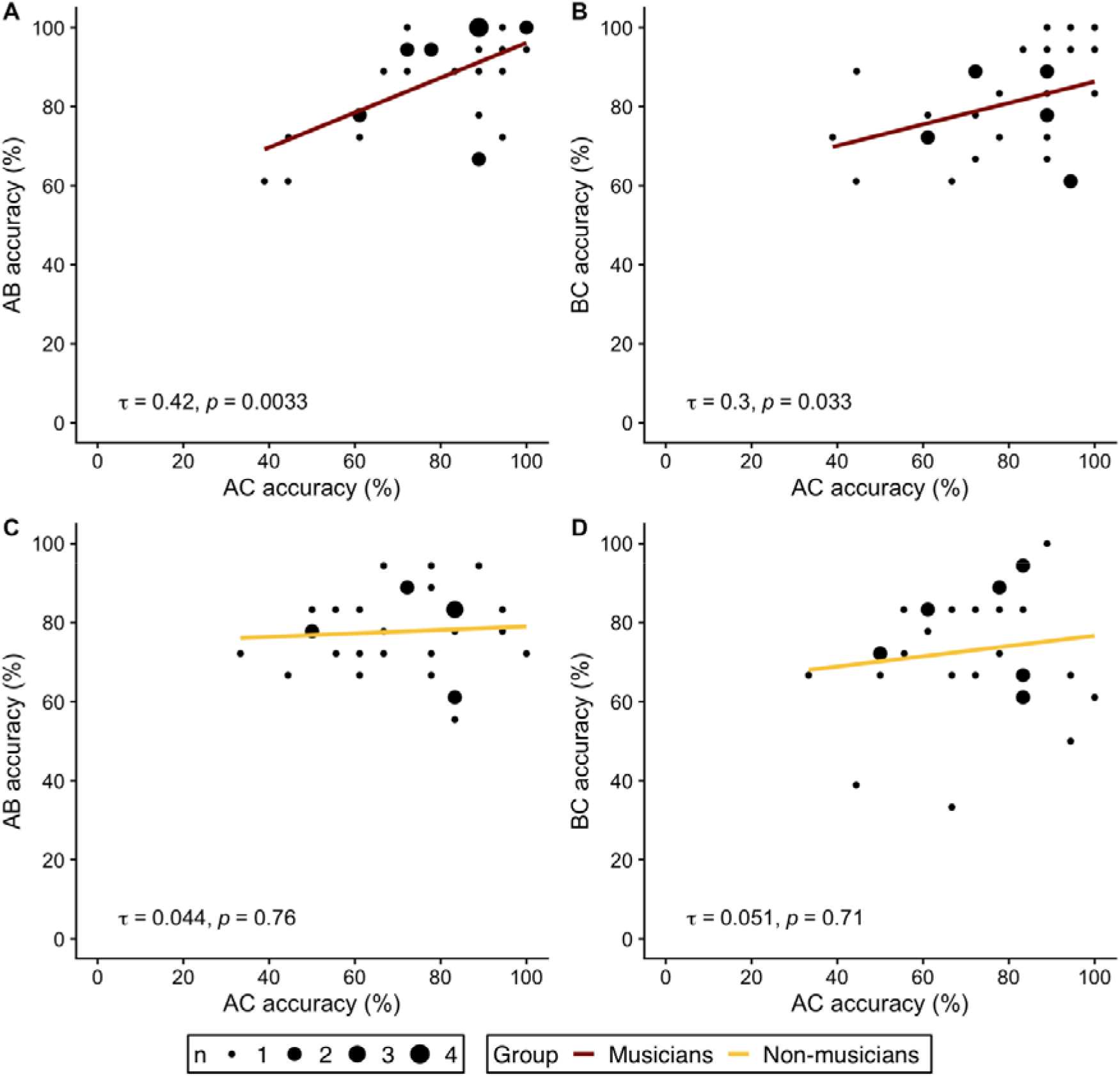
Correlations of indirect trials (AC accuracy on the respective x axis) and direct trials (AB, BC accuracy on the respective y axis) in musicians (A & B, red) and non-musicians (C & D, yellow). (A, C) Correlation of AC- and AB-trials. (B, D) Correlation of AC- and BC-trials. τ refers to the correlation coefficient from Kendall’s τ. Dot size represent the number of identical values.

### Correlation of direct and indirect trials with music-related variables

We reasoned that if professional musicianship shifts memory integration strategies from a mainly encoding-based strategy to a more retrieval-based strategy, we should expect distinct correlations between trial type parameters and music-related variables. We therefore examined relationships between indices of musical activity with accuracy and RTs in indirect and direct trials. In non-musicians, we found no correlations between any variable of musical activity (i.e. MBEA Scale performance, music listening and concert visits) and accuracy or RTs in both trial types (Supplementary Table 2). In musicians, MBEA scale performance and concert visits were not associated with accuracy or RTs of both trial types (Supplementary Table 2). Music listening however correlated with accuracy in indirect trials (τ = −0.33, p = 0.026), thus suggesting that musicians who listen to music for a longer duration per day have a lower performance in answering AC-trials (see Figure 5A).

**Figure 5.**
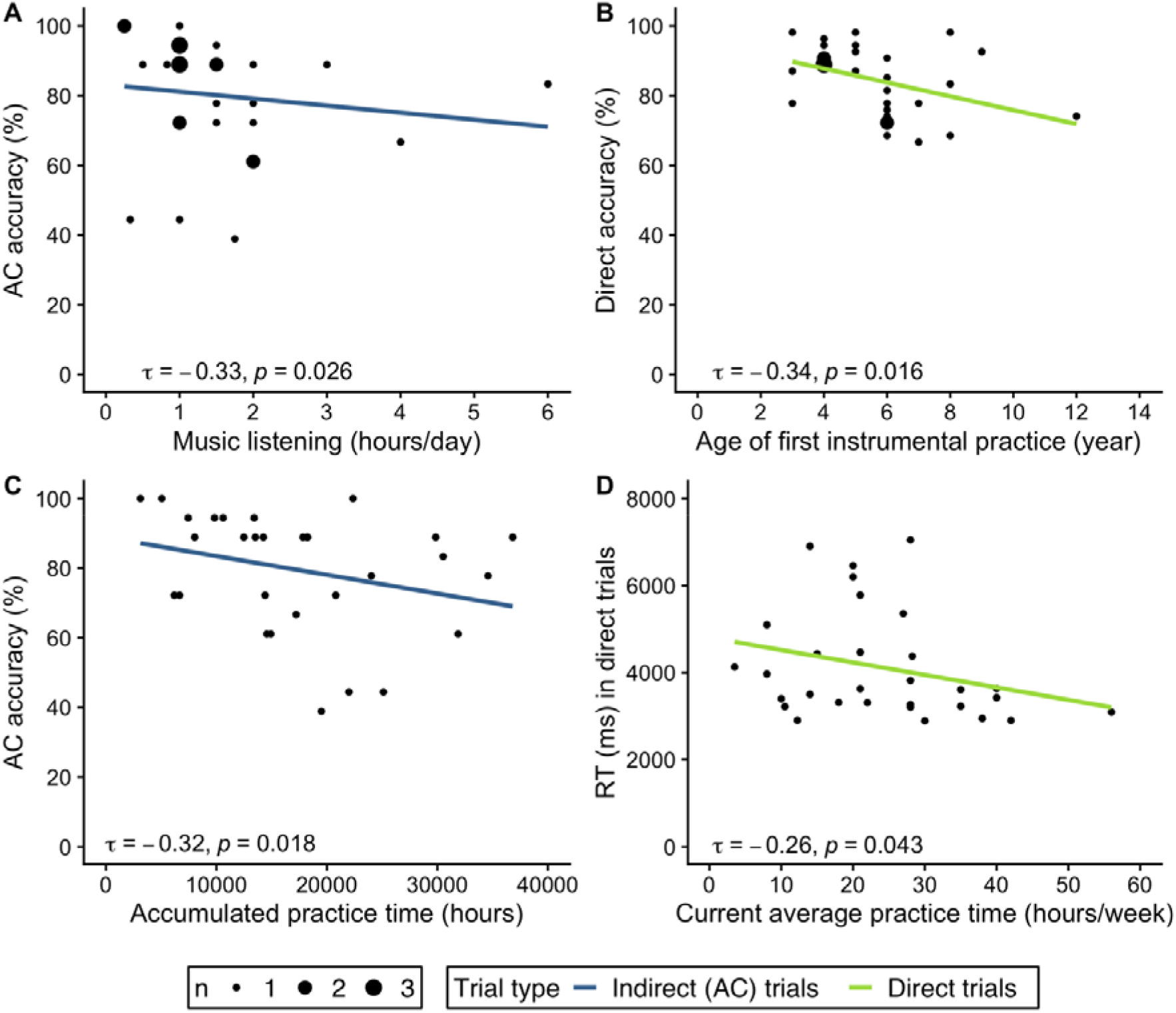
Correlations of music-related variables with accuracy and RTs in musicians. (A) Correlation of the average hours of daily music listening during the last 12 months with accuracy in AC-trials. (B) Correlation of the age of first instrumental practice with accuracy in direct trials. (C) Correlation of the average practice time per week during the last 12 months with RTs in direct trials. (D) Correlation of the accumulated practice time with accuracy in AC-trials. τ refers to the correlation coefficient from Kendall’s τ. Dot size represents the number of identical values.

We additionally examined correlations between accuracy and RTs with variables of active music making in musicians. Age of first instrumental practice correlated with accuracy of direct trials (τ = −0.34, p = 0.016). Musicians with a younger age of first instrumental practice showed higher performance in direct trials (Figure 5B). No correlations of the age of first instrumental practice with indirect trials or RTs were observed. Years of instrument playing was not associated with accuracy or RTs of both trial types.

Accumulated practice time was not associated with direct accuracy or RTs in any trial type. However, there was a significant negative correlation of accumulated practice time and indirect accuracy (τ = −0.32, p = 0.018). Musicians with higher accumulated hours of practice had lower performance in indirect trials (see Figure 5C). We reasoned that accumulated practice time is also confounded with age, with older persons having higher values of cumulative practice times. Accordingly, our data showed a significant correlation of age and cumulative practice time (τ = 0.32, p = 0.014). We therefore calculated a partial correlation between indirect accuracy and accumulated practice time and controlled for age. Results of the partial correlation analysis were still significant (τ = −0.28, p = 0.031), indicating that age did not account for the negative bivariate correlation of indirect accuracy and accumulated practice time.

Finally, we analyzed associations of practice times with accuracy and RTs. Current practice time (i.e. the average practice time per week during the last 12 months) correlated with RTs in direct trials (τ = −0.26, p = 0.043), suggesting that musicians with higher current practice times reacted faster in direct trials (see Figure 5D). Current practice time did not correlate with RTs in indirect trials or accuracy of direct or indirect trials. General average practice time (i.e. average practice time per week across all decades of musical activity) did not correlate with accuracy or RTs in any trial type.

## Discussion

We investigated how musicians and non-musicians use melodies to build associations with non-musical information and form integrated memory representations. Using an associative inference task with visual and musical stimuli, we compared accuracy and RTs of professional musicians and non-musicians for memory of direct visual-melodic associations as well as for indirect inferential associations in which melodies link otherwise unrelated visual information. Although musicians outperformed non-musicians in both trial types, non-musicians were also able to use melodies to build associations with non-musical information. Our findings however suggest that musicians and non-musicians use strikingly different strategies for memory integration.

Accuracy of both musicians and non-musicians was above chance level in direct and indirect trials. This indicates that our participants could reliably memorize and retrieve not only associations of objects and melodies, but could also use melodies to bind distinct and previously unrelated visual information into an integrated memory representation. Importantly, these results were found not only in musically trained individuals but also in persons without any musical training. It has previously been proposed that music can easily be associated with non-musical information, such as autobiographic or semantic memories^6,8,9^. Both everyday experience and previous research show that music can be a powerful stimulus for memory retrieval. For instance, musical cues were used in a study on the neural correlates of autobiographic memories of different levels of specificity^36^. The results showed that familiar music can serve as cue for a wide range of autobiographical memory episodes, even if these episodes had only rarely been retrieved from memory before. The relationship between music, involuntary musical imagery (i.e. earworms) and memory for associated events was investigated in a recent study^37^, in which participants were first exposed to soundtracks that were likely to evoke involuntary musical imagery. In a second session, the soundtracks were paired with unfamiliar cartoon movies. After different delay periods, participants were cued with these soundtracks to remember details of the related cartoons. The results showed that across delays of 1-2 weeks, participants could not only recognize the soundtracks but also that music could evoke associated event information previously presented in the movies. Memory of music and associated event knowledge was further predicted by the amount of mental replay of the music during the delay. Importantly, the latter study included non-musicians or participants with a few years of formal musical instrument training, but no professional musicians^37^. These results as well as the findings from our study support the hypothesis that the process of building links between music and non-musical memories can happen with surprising efficacy, is not confined to musically trained persons and extends well beyond memory of salient autobiographical episodes.

Not surprisingly, musicians have been found to have superior auditory memory compared to non-musicians, not only for musical but also for non-musical auditory stimuli^38^. In both musicians and non-musicians, however, auditory memory was inferior to visual memory, which was comparable between groups^38^. Similarly, a meta-analysis found that, compared to non-musicians, musicians have a better performance in memory tasks, with a small effect for long-term memory and medium effect sizes for short-term and working memory tasks^13^. Better memory performance was however dependent on stimulus type. For short-term and working memory tasks, the memory advantage of musicians was large for tonal stimuli, moderate for verbal stimuli and small or null when visuospatial stimuli were involved. In a more recent study, visual and auditory short-term memory in musicians and non-musicians was compared using different categories of stimuli (i.e. verbal, non-verbal with contour, non-verbal without contour)^39^. Stimulus sequences with contour included up and down variations based on loudness (auditory condition) or luminance (visual condition). Musicians selectively performed better in both visual and auditory contour and auditory non-contour conditions, whereas memory performance in verbal conditions was comparable. These results suggest that musical activity preferentially trains memory domains that are closely related to musical skills. In line with this, research on other fields of expertise such as chess, medicine or mental calculations revealed that experts mainly have a domain-specific memory advantage for meaningful information within their field of expertise^40,41^. It seems therefore likely that performance differences across our two groups are mainly driven by superior auditory memory in musicians rather than by an overall higher level of memory performance.

To correctly perform in AC-trials, A- and C-stimuli that were not experienced together have to be linked at some point between encoding and retrieval. As outlined previously^18,19,21,26,42^, two complimentary processes have been postulated that may support memory integration. First, memory integration may be achieved by an integrative encoding mechanism^18,19,26,43^. During encoding of BC-pairs, previously studied AB-pairs become reactivated via the overlapping B-stimulus. Thus, integrated ABC-representations are formed already during encoding and are readily available for later inferential AC-decisions, since the underlying AB- and BC-pairs do not have to be retrieved separately^26^. In fact, a previous study revealed that response times for untrained inferential associations could be as fast as for trained direct associations, lending support for the idea that integrated memories can already be constructed during encoding^43^. Second, integration of distinct but related memories can occur during retrieval. In this case, individual AB- and BC-pairs have to be remembered separately and are finally recombined for AC-decisions. This process has been termed recombination at retrieval^18^ and is more flexible, but results in slower responses, since additional cognitive processes are necessary at retrieval that are not required for retrieving direct pairs^42^. Neuroimaging studies revealed that processes in the hippocampus support memory integration both during encoding and retrieval^19,26,44–47^. In line with these neuroimaging results, patients with lesions in the hippocampus and MTL were found to have deficits in memory integration that cannot solely be explained by difficulties in associative memory^21^ and show reduced performance in making inferences between items of overlapping memory networks^48^.

Our data suggest that musicians and non-musicians used qualitatively different strategies for memory integration. Non-musicians showed faster responses in correct AC-trials compared to correct direct trials. We therefore assume that non-musicians use an integrative encoding strategy in which they build a melodic link between A- and C-stimuli, i.e. an ABC-triplet that is formed when the BC-pair is presented. Thus, an integrated representation is already available at retrieval for AC-decisions.

However, the studied direct pairs require additional processing time. Notably, the direct pairs involved melodies, which might be more difficult and need more processing time to recognize for non-musicians than purely visual information (i.e. AC-trials). In line with this, previous research has shown that auditory memory for a wide range of auditory stimuli is inferior to visual memory in both musically trained and untrained individuals^38,49^. The analysis of accuracy in corresponding AB-, BC- and AC-trials of one triplet as well as the correlation analysis of accuracy further support the idea of different strategies in the two groups. Since non-musicians presumably memorize A-and C-stimuli together already during encoding, they are less dependent on knowledge of underlying AB- and BC-pairs for AC-decisions. Therefore, non-musicians can still correctly decide on AC-pairs even if they do not correctly remember the corresponding AB-and/or BC-pairs or reconstruct AB- and BC-pairs from knowledge of AC-pairs. By contrast, musicians seem to base their AC-decisions mainly on memory of the underlying AB-and BC-pairs, which they recombine flexibly at retrieval in AC-trials.

Our correlation analyses showed that an earlier start of instrumental practice was associated with higher performance in direct trials. In previous studies, an earlier start of musical training was associated with changes in auditory and motor brain areas^50–52^ and higher accuracy in a visual memory task^53^. Compared to musicians with a later start of musical training, early-trained musicians were found to have higher temporal precision in piano playing^54^ and a better performance in music-related tasks, such as simple melody discrimination^55^, auditory-motor and visual-motor rhythm synchronization tasks^56–58^. In our study, the age of onset of musical training was also associated with higher performance in memory of object-melody associations, but not with accuracy in inferential object-object associations. Performance in direct accuracy may be linked to the ability to discriminate and memorize melodies over a longer time period, which might be more developed in early-trained musicians. Memory of inferential object-object associations seems to be less influenced by musical training and is presumably supported by the basic integrative encoding mechanism that seems to be available to most people, irrespective of musical activity. However, performance in direct trials not solely reflects onset of musical training but seems to be modified by practice factors, since current musical practice times were associated with faster responses in direct trials in the musician group. This finding is in line with a previous study reporting correlations between daily instrumental practice and reaction times in visual memory tasks that could reflect better attentional processes in musicians with higher practicing times^53^.

Surprisingly, a higher performance in AC-trials was associated with both a shorter average duration of music listening per day and with lower accumulated practice times. It is conceivable that passive listening to music for long durations is not helpful for memorizing melodies. Instead, for memorizing melodies and object-melody pairs, it might be more important to have a rather analytical and focused style of music listening. In general, attentional focus is closely related to learning and memory^59^. For instance, attention modulates hippocampal activity patterns with stronger representations of the correct attentional state during encoding of items that were remembered subsequently^60^. Attentional focus during music listening might also influence the experience of music, related autobiographical memories and activation of brain networks^61^. Similarly, although accumulated practice time has been suggested as a modulator of neuroplasticity related to musical activity^62^, it might not only be the amount but also the nature of practice that determines musical proficiency^40^. Similarly important for development of superior performance skills is deliberate practice, i.e. practice situations, in which the focus is on improving a particular aspect and which allow for repetitions and direct feedback^40,63^. That way attentional focus seems to be an important aspect of musical training and might also contribute to associations between measures of musical training intensity and task performance.

Compared to non-musicians, musical information has a higher relevance and is more closely related to personal behavior in professional musicians, who are often required to select relevant melodies and memorize them actively and consciously. For musicians, music must not only be recognized but also reliably recalled and imitated. This is especially relevant for musical improvisation, but also for performances without sheet music. It has previously been proposed that memorizing melodies mostly involves chunking and consolidation of small musical ordered segments and that musical training may foster acquisition of controlled and active learning strategies (e.g. chunking)^13^. In our study, such an active learning strategy might have contributed to task performance, so that musicians could memorize and recombine the underlying chunks, i.e. pairs of melodies and objects. We therefore assume that musicians can not only rely on a default integrative encoding mechanism, but additionally have access to a second strategy (i.e. recombination at retrieval), allowing them to use melodies for associations with non-musical information in a more deliberate and flexible way. Since non-musicians are less trained to actively recall and reproduce melodies, they are prone to rely on a recognition-based mechanism of integrative encoding.

On a general level, our findings show that music can function as an efficient mnemonic device even for biographically irrelevant non-musical information. Musical mnemonics are already used in pedagogical settings to teach contents of different complexity in an easy and enjoyable way. A popular example is the Schoolhouse Rock, a series of short educational musical animated videos that were broadcasted between TV children’s shows between the 70s and 80s and presented a variety of different topics from mathematics to grammar to science in songs^64^. Musical mnemonics have also been used for education in academic settings for more complex information such as health science^65^ or biochemistry and molecular biology^66^. The reported application of musical mnemonics in education does however not necessarily prove that information presented in music is indeed more memorable. Therefore, previous research also evaluated the potential of music to enhance memory of non-musical information and yielded inconclusive results. While some studies did not find any differences in memory for sung and spoken information in healthy participants^67–69^, other studies found an positive effect of musical mnemonics with better recall of sung compared to spoken words and lyrics^70–72^. The capacity of music to improve verbal memory has also been investigated in patients with memory impairments. Compared to presenting verbal information in spoken form, sung material improved word memory in patients with multiple sclerosis^73^ as well as learning and recall of verbal narrative stories in stroke patients 6 months post-stroke^74^ and enhanced recognition of sung lyrics in patients with Alzheimer’s disease^75^. Our findings suggest that melodies might also be an effective mnemonic for visual information and that music can be used to form memory networks of related memory episodes.

How this finding might contribute or be applied in pedagogical or therapeutical settings needs to be addressed in future studies.

Taken together, the findings reported here suggest that both musicians and non-musicians can use melodies to efficiently memorize and retrieve visual information. However, musically trained and untrained individuals differ substantially in how they use melodies for memory integration. Our results suggest that integrative encoding is a default mechanism for integration of musical and non-musical stimuli that is available to a surprising degree even to musically untrained subjects. We speculate that this more passive and recognition-based mechanism may reflect a basic ability to intuitively attach sounds to objects with no or little conscious effort. We cannot be sure whether this is specific to music, but it appears likely that integrative encoding may contribute significantly to the everyday experience of music-evoked episodic memories. By contrast, recombination at retrieval seems to be a more active and recall-based strategy for memory integration that apparently depends on an expert ability to maintain and discriminate musical stimuli across memory delays. Future studies should investigate if distinct behavioral strategies in musicians and non-musicians are underpinned by differential neural correlates or if task performance in musicians is related to activation in networks that are associated with their own instrumental practice. Moreover, it will be important to investigate how visual-melodic memory integration persists across extended memory delays and how integrative encoding can be used to facilitate the learning of new information in normal subjects and subjects with learning impairments.

## Data availability statement

Anonymized data of the study are available at the Open Science Framework and can be accessed at https://osf.io/63mep/

## Acknowledgements

We thank Bernd Enders for the support in selecting and generating the musical stimuli. We also thank all study participants for supporting our study.

## Author contributions

All authors designed and conceptualized the study. M.H. collected the data. All authors analyzed and interpreted the data. All authors drafted, reviewed and edited the manuscript.

## Supplementary information

**Supplementary Table 1.** List of musical stimuli.

| Composer | Title | Genre | Length |
| --- | --- | --- | --- |
| Komarowski, Anatoli | Violin concerto | Classic | 00:07 |
| Badings, Henk | Scherzo Pastorale (from Little Piano Pieces) | Classic | 00:06 |
| Bartok, Bela | Bulgarian Rhythms - 4th mvt | Classic | 00:06 |
| Thielemans, Toots | Bluesette | Jazz | 00:07 |
| Bolling, Claude | Javanaise (from Suite for Flute and Jazz Piano) | Jazz/Classic | 00:06 |
| Bortkiewicz, Sergei E. | The Princess and the Pie (from Andersen's Fairy Tales) | Classic | 00:05 |
| Debussy, Claude | Passepied (from Suite Bergamasque) | Classic | 00:07 |
| Chabrier, Emmanuel | Espana | Classic | 00:08 |
| Gulda, Friedrich | Moderato (from Play Piano Play) | Jazz | 00:07 |
| Busch, Elliot | Ivory Rag | Jazz | 00:08 |
| Janáček, Leoš | No. 5 (from On an Overgrown Path) | Classic | 00:05 |
| Janáček, Leoš | Moderato (from Sinfonietta) | Classic | 00:07 |
| Khachaturian, Aram | Andantino (from Album for Children No. 1) | Classic | 00:10 |
| Manuel, Horacio | Balada Da Felicidade | - | 00:08 |
| Mays, Lyle | Chorinho | Jazz | 00:06 |
| Pachelbel, Johann | Magnificat fugue secundi toni No. 2 | Classic | 00:06 |
| Poulenc, Francis | Mouvements perpétuels | Classic | 00:06 |
| Prokofiev, Sergei | Vivace (from Piano Sonata No. 6) | Classic | 00:05 |
| Skrjabin, Alexander | No. 4 Lento (from 5 Préludes) | Classic | 00:06 |
| Portillo de la Luz, César | Son al Son | Traditional | 00:06 |
| Taki, Rentaro | Moon over ruined castle | Traditional/Folk song | 00:07 |
| Troup, Bobby | Their hearts were full of spring | Pop | 00:05 |
| Tchaikovsky, Pyotr I. | Chanson Triste op. 40/No. 2 | Classic | 00:06 |
| Joplin, Scott | Fig Leaf Rag | Jazz | 00:07 |
| Gounod, Charles | Funeral March of a Marionette | Classic | 00:08 |
| Moszkowski, Moritz | Spanish Dance No. 2 | Classic | 00:05 |
| Copland, Aaron | Three Moods, Jazzy | Jazz | 00:09 |
| O'Hagan, Jack | Road to Gundagai | Traditional/Folk song | 00:08 |
| Gershwin, George | Rialto Ripple Rag | Classic/Jazz | 00:07 |
| Ascher, Emil | Arabian Wedding March | Classic | 00:08 |
| Riedel, Georg | Idas Sommarvisa | Folk song | 00:09 |
| Bonnet, Carlos | La partida (Vals venezolana) | Traditional | 00:07 |
| - | Saka Naka | Traditional/Folk song | 00:08 |
| - | Klezmeron | Traditional | 00:08 |
| Johnson, Laurie | The Avengers | Theme Song/Film song | 00:07 |
| Stein, Herman | Gunsmoke | Theme Song/Film song | 00:08 |
| Vars, Henry | Flipper | Theme Song/Film song | 00:07 |
| Rose, David | Little House -A new beginning | Theme Song/Film song | 00:06 |
| Beethoven, Ludwig v. | Return to Ulster (from 25 Irish Songs) | Classic | 00:10 |
| Tugarinov, J. | Vo pole berjesa stojala | Traditional/Folk song | 00:10 |
| Anonymous/Thomas Oliphant | The Ash Grove | Traditional/Folk song | 00:07 |
| Copland, Aaron | Appalachian Suite | Classic | 00:08 |

**Supplementary Table 2.** Correlations between musical activity variables, accuracy and RTs in indirect and direct trials in musicians and non-musicians

| Group | Variable | Accuracy |  | RTs |  |
| --- | --- | --- | --- | --- | --- |
|  |  | Indirect | Direct | Indirect | Direct |
| Non-musicians | MBEA Scale | 0.16 (0.243) | 0.11 (0.445) | -0.13 (0.329) | -0.07 (0.588) |
|  | Music listening | 0.16 (0.253) | 0.10 (0.45) | 0.17 (0.209) | 0.19 (0.151) |
|  | Concert visits | -0.22 (0.116) | -0.06 (0.662) | 0.02 (0.871) | 0.05 (0.704) |
| Musicians | MBEA Scale | -0.11 (0.429) | -0.09 (0.493) | -0.01 (0.971) | -0.02 (0.912) |
|  | Music listening | -0.33* (0.026) | -0.20 (0.158) | 0.05 (0.699) | 0.13 (0.354) |
|  | Concert visits | -0.17 (0.238) | -0.18 (0.189) | 0.07 (0.609) | -0.002 (0.985) |
|  | Age of first instrumental practice | -0.20 (0.157) | -0.34* (0.016) | -0.13 (0.332) | 0.09 (0.522) |
|  | Years of instrument playing | -0.13 (0.361) | -0.09 (0.472) | 0.18 (0.162) | 0.12 (0.351) |
|  | Accumulated practice time | -0.32 (0.018)* | -0.22 (0.099) | 0.19 (0.135) | -0.05 (0.697) |
|  | General average practice time<sup>a</sup> | -0.24 (0.081) | -0.24 (0.071) | 0.10 (0.432) | -0.17 (0.175) |
|  | Current average practice time<sup>b</sup> | -0.08 (0.547) | -0.06 (0.667) | -0.18 (0.179) | -0.26* (0.043) |

Correlation coefficients refer to Kendall's $\tau$ . Values in parentheses are p values of the respective correlation. \* denotes $p < 0.05$ ; <sup>a</sup> Average practice time per week across all decades of musical activity; <sup>b</sup> Average practice time per week during the last 12 months.

